# Drug Screening Platform Using Human Induced Pluripotent Stem Cell-Derived Atrial Cardiomyocytes and Optical Mapping

**DOI:** 10.1101/2020.07.14.203232

**Authors:** Marvin G. Gunawan, Sarabjit S. Sangha, Sanam Shafaattalab, Eric Lin, Danielle A. Heims-Waldron, Vassilios J. Bezzerides, Zachary Laksman, Glen F. Tibbits

**Affiliations:** Molecular Cardiac Physiology Group, Departments of Biomedical Physiology and Kinesiology and Molecular Biology and Biochemistry, Simon Fraser University, 8888 University Drive, Burnaby, BC, V5A 1A6, Canada; Tibbits Research Team, Cellular and Regenerative Medicine Centre, BC Children’s Hospital Research Institute, 950 West 28th Avenue, Vancouver, BC, V5Z 4H4, Canada; Division of Cardiology, Faculty of Medicine, University of British Columbia, 317 – 2194 Health Sciences Mall, Vancouver, BC, V6T 1Z2, Canada; Centre for Heart and Lung Innovation, St. Paul’s Hospital, 1081 Burrard St, Vancouver, BC, V6Z 1Y6, Canada; Department of Cardiology, Boston Children’s Hospital, 300 Longwood Ave., Boston, MA, 02115, USA

**Author notes:** These authors contributed equally to this article (Marvin G. Gunawan, Sarabjit S. Sangha, Sanam Shafaattalab). Corresponding author Correspondence Dr. Glen F. Tibbits, Room 2084, British Columbia Children’s Hospital Research Institute, 950 West 28^th^ Avenue, Vancouver, British Columbia, V5Z 4H4, Canada. Author Contributions Marvin G. Gunawan and Sarabjit S. Sangha: conception and design, collection and assembly of data, data analysis and interpretation, manuscript writing; Sanam Shafaattalab: experimental design support, data interpretation, manuscript writing; Eric Lin: designed and built the optical mapping system (hardware and software); Danielle A Heims-Waldron: cell culture, Vassilios J. Bezzerides: data collection, data analysis and interpretation, manuscript writing; Zachary Laksman and Glen F. Tibbits: conception of study, manuscript writing support and review, data interpretation, financial support.

**Keywords:** drug screening, atrial fibrillation, cardiomyocyte subtype, atrial differentiation, human induced pluripotent stem cells

## Abstract

Current drug development efforts for the treatment of atrial fibrillation (AF) are hampered by the fact that many preclinical models have been unsuccessful in reproducing human cardiac atrial physiology and its response to medications. In this study, we demonstrated an approach using human induced pluripotent stem cell-derived atrial and ventricular cardiomyocytes (hiPSC-aCMs and hiPSC-vCMs, respectively) coupled with a sophisticated optical mapping system for drug screening of atrial-selective compounds *in vitro*.

We optimized differentiation of hiPSC-aCMs by modulating the WNT and retinoid signalling pathways. Characterization of the transcriptome and proteome revealed that retinoic acid pushes the differentiation process into the atrial lineage and generated hiPSC-aCMs. Functional characterization using optical mapping showed that hiPSC-aCMs have shorter action potential durations and faster Ca^2+^ handling dynamics compared to hiPSC-vCMs. Furthermore, pharmacological investigation of hiPSC-aCMs captured atrial-selective effects by displaying greater sensitivity to atrial-selective compounds 4-aminopyridine, AVE0118, UCL1684, and vernakalant when compared to hiPSC-vCMs.

These results established that a model system incorporating hiPSC-aCMs combined with optical mapping is well-suited for pre-clinical drug screening of novel and targeted atrial selective compounds.

## Introduction

The advent of human induced pluripotent stem cell-derived cardiomyocytes (hiPSC-CMs) has revolutionized the field of cardiac research. It has enabled the study of cardiac diseases in a patient-specific and human-relevant *in vitro* model system which provides a unique opportunity for clinical translation^1^. Furthermore, the ability to differentiate chamber-specific cardiomyocytes allows for a more precise study of cardiac disease physiology and pharmacology.

The cardiomyocytes of the lower (ventricles) and upper (atria) chambers have distinct characteristics that arise from differential developmental pathways. Previous work *in vivo* has shown that the expression patterns of retinoic acid and retinaldehyde dehydrogenase 2 (RALDH2) are important determinants of the atrial fate^2–5^. These results were later recapitulated in a pivotal study by Lee & Protze et al.^6^ who determined that atrial cardiomyocytes (aCMs) differentiated from human embryonic stem cells (hESCs) originate from a unique mesoderm characterized by robust RALDH2 expression. This study established an atrial differentiation protocol that included the addition of retinoic acid. Retinoic acid has also been utilized to selectively differentiate hESCs and hiPSCs into aCMs in other studies^6–10^.

The distinct properties of the atrial and ventricular cardiomyocytes are determined by the differential expression of unique sets of ion channels and other proteins that optimize their specific function. Drugs that target atrial ion channels selectively can therefore produce differences in pharmacological function in the two chambers. This atrial-selective pharmacology is of utmost interest in the study and treatment of atrial-specific diseases such atrial fibrillation (AF), which is the most common heart rhythm disorder. Investigating atrial-selective pharmacology can assist and guide novel cardiac drug development as well as improving both safety and efficacy by avoiding potential toxic electrophysiologic effects on the ventricular chambers.

The differential pharmacology of stem cell-derived aCMs was studied previously by Laksman et al.^7^ who showed that flecainide can rescue the AF phenotype in a dish. Other studies have also studied the selective pharmacological effects of agents on hiPSC-derived aCMs but have largely focused on being proof-of-concept studies using limited number of test compounds and standard measurement systems that are low in throughput^9,10^. With a focus on translation, a pre-clinical model platform that characterizes pharmacological activity must capture the main cardiac functional signatures that most closely mimic and predict human cardiac physiology and drug responses. As such, we established in this study an *in vitro* assay platform by combining hiPSC-derived atrial cardiomyocytes (hiPSC-aCMs) and high-content optical mapping, a non-invasive all-optical system that simultaneously measures membrane potential (V_m_) and Ca2+ transients at a high-resolution in a monolayer tissue format.

We first demonstrate a selective hiPSC-aCM differentiation protocol by modifying the well characterized GiWi protocol^11^ through the controlled introduction of retinoic acid. The recapitulation of the human atrial phenotype of the hiPSC-aCMs was validated with assays that measure the expression of gene transcripts and proteins, as well as functional signatures. We then demonstrate the utility of our platform as an atrial-selective drug screening tool by using existing clinical and experimental drugs. The model established in this study adds to our current understanding of the utility of stem cell derived cardiomyocytes in pre-clinical and translational research focused on screening new pharmacological agents.

## Methods and MATERIALS

A detailed Methods section is available in the Supplemental Information.

### Maintenance and Expansion of hiPSCs

hiPSCs (WiCell, IMR90-1) were maintained and expanded in mTeSR1 medium and feeder-free culture using 6-well plates coated with Matrigel. Using Versene (EDTA), hiPSCs were passaged every 4 days or ∼85% confluency at 1:15 ratio. Passaged hiPSCs were cultured with mTeSR1 supplemented with 10 µM Y27632 for the first 24 hours and the mTeSR1 was exchanged daily during cell culture maintenance.

### Directed Differentiation of hiPSCs into Atrial and Ventricular Subtypes

hiPSC-derived ventricular cardiomyocytes were differentiated by employing a modified GiWi protocol^11^ that we previously described^12^. In brief, hiPSCs were seeded at a density of 87,500 cells/cm^2^. At day 0, differentiation was initiated using 12 µM CHIR99021. At day 3, the cells were incubated with 5 µM IWP-4. At day 5, the media were refreshed with RPMI-1640 supplemented with B27 minus insulin. At day 7, the medium was replaced with cardiomyocyte maintenance media (RPMI-1640 supplemented with B27 with insulin). Thereafter, cardiomyocyte maintenance medium was replaced every 4 days. For the atrial differentiation protocol, retinoic acid (RA) addition was first optimized in pilot studies (Figure S2 & S3) and determined to be 0.75 µM RA every 24 hours from days 4-6.

### Flow Cytometry

hiPSC-aCMs and hiPSC-vCMs at Day 20-30 post-differentiation were dissociated into single cells as described in the Supplemental Information. The harvested cells were fixed in 4.1% PFA solution for 25 min and then washed and permeabilized in Saponin/FBS. Cells were subsequently incubated overnight in primary mouse-cTnT (1:2000) and rabbit-MLC2V (1:1000) antibodies. Subsequently, the cells were washed and incubated for one hour in secondary goat anti mouse Alexa-488 (1:500) and goat anti rabbit Alexa-647 (1:2000) antibodies, respectively. Cells were then washed and suspended in PBS for analysis. All analyses were performed using the BDJAZZ Fluorescence Activated Cell Sorter.

### mRNA Expression Profiling

Gene expression profiling was conducted using multiplexed NanoString and real time quantitative PCR (qPCR). Pooled total RNA was used in both assays. The extracted RNA was reverse transcribed into cDNA which was used in the qPCR assay. Oligonucleotide sequences are described in Table S7. The multiplexed mRNA profiling was conducted using a NanoString Technologies (Seattle, WA) platform with a custom code set containing 250 gene probes. Analysis was performed on the Nanostring Sprint instrument and nSolver analysis software with the Advanced Analysis module.

### Atrial Natriuretic Peptide Measurement

The levels of atrial natriuretic peptide (ANP) of hiPSC-aCMs and -vCMs were measured by a competitive enzyme-linked immunosorbent assay (ELISA) using a commercially available kit (Invitrogen, CA). The assay was conducted according to the manufacturer’s protocol and was measured using a spectrophotometric plate reader.

### Cardiomyocyte Enrichment

For cardiac enrichment, hiPSC-aCMs and -vCMs at day 20-30 post-differentiation were dissociated into single cells which were then enriched using a MidiMACS PSC-derived Cardiomyocyte Isolation Kit (Miltenyi Biotec, Germany) according to the manufacturer’s protocol. Enriched hiPSC-CMs were seeded on Matrigel-coated 24-well plates at a seeding density of 600,000 cells per well.

### Patch-clamp Recordings

Single hiPSC-aCMs and -vCMs were plated on gelatin (0.1%) and Geltrex (1:10) at 30,000 cells per well. After 48 hours in culture, glass electrodes were used to achieve the whole-cell configuration with single hiPSC-CMs and only cells with gigaohm seals were used for further analysis. The formulation for internal and external recordings solutions are outlined in the Supplemental Information. Current recordings were performed using an Axon Instruments 700B amplifier and digitized at 20 KHz. All recordings were performed at 33-35 °C as maintained. For pacing at 1 Hz, gradually increasing amounts of current were injected with a 1 ms pulse width until reliable action potentials (APs) were triggered. The maximal upstroke velocity was determined by calculating the maximum derivative and the resting membrane potential was measured during a 5 second epoch without spontaneous activity one minute after break-in. Further details on data analysis are found in the Supplemental Information.

### Optical Mapping

Optical mapping recordings were performed on enriched monolayers of hiPSC-aCMs and -vCMs cultured in a 24-well plate format at Day 45-60 post-differentiation. Imaging experiments were conducted using Ca^2+^ Tyrode’s solution (formulation found in Supplemental Information). The hiPSC-CMs were loaded with RH-237, blebbistatin, and Rhod-2AM sequentially before imaging as described^12,13^. Both RH-237 and Rhod-2AM were excited by 530 nm LEDs. Images were acquired at a frame rate of 100 frames/second by a sCMOS camera (Orca Flash 4.0 V2, Hamamatsu Photonics, Japan) equipped with an optical splitter. The cells were paced using programmable stimulation. Data collection, image processing, and initial data analysis were accomplished using custom software. The multi-well optical mapping system was custom engineered in the lab based on a system as described previously^17^. Further details are found in the Supplemental Information.

### Pharmacological Analyses

The drugs used in this study are listed Table S8. Drug stocks were further diluted in Ca^2+^ Tyrode’s solution prior to pharmacological testing with the final DMSO concentration in the experimental solution not exceeding 0.03% (v/v). Drug effects were studied in serum-free conditions (i.e. Ca^2+^ Tyrode’s and drug only) at four doses by sequentially increasing the drug concentration in the same well with recordings at 20-minute intervals.

### Statistical Analysis

Further details on data and statistical analysis can be found in the Supplemental Information. Unpaired t-tests were conducted to compare two groups (i.e. hiPSC-aCMs vs. hiPSC-vCMs) in the analysis of qPCR, ELISA, patch clamp recordings, and optical mapping (baseline condition and normalized drug effects). Analysis of dose-dependent effects were performed using one-way ANOVA and Dunnett’s post-hoc test. All data are presented as mean ± SEM unless noted otherwise. Significance level for all statistical analysis was set at p < 0.05 with the following notation: *p < 0.05, **p < 0.01, ***p < 0.001.

## Results

### RA Treatment Drives Cardiac Differentiation into Atrial Phenotype

We first optimized the atrial differentiation protocol by altering the concentration and timing of retinoic acid (RA) based on the molecular signatures of an atrial phenotype as measured by qPCR and flow cytometry (Figures S2 & S3). Higher dose of RA reduced cardiac differentiation efficacy defined by the decrease in the cTnT^+^ proportion of the total cell population as measured by flow cytometry (Figure S2A). The finalized protocol to generate hiPSC-aCMs included the addition of RA at 0.75 µM every 24 hours on days 4, 5, and 6 (Figure 1A) which was found as a balance between sufficiently driving atrial differentiation as defined by decreased ventricular marker myosin light chain 2 – ventricular paralog (MLC-2v) while having no impact cardiac differentiation efficacy (Figure S2 & S3).

**Figure 1:**
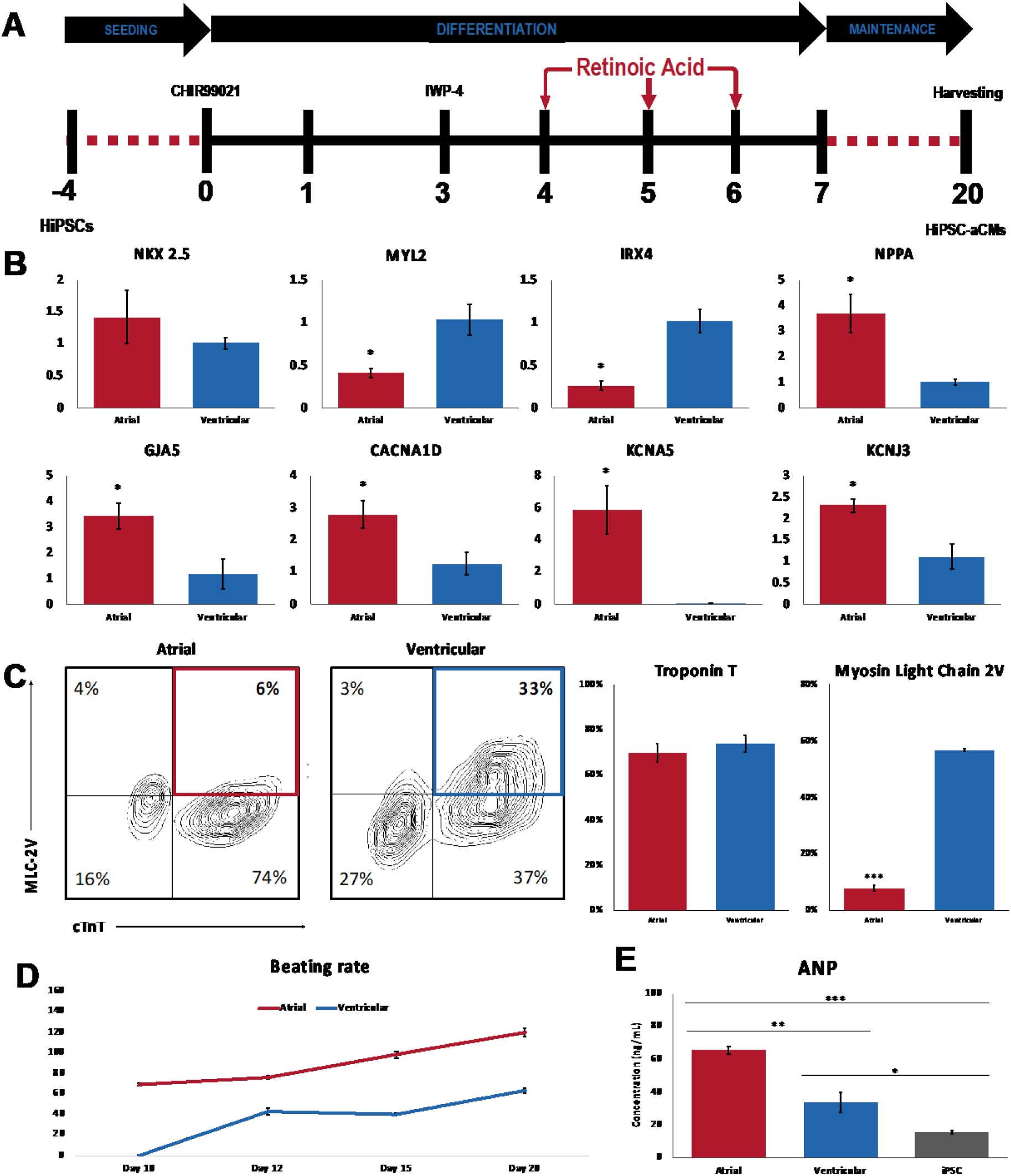
Directed differentiation of hiPSC-derived atrial and ventricular CMs. A) Schematic depicting the atrial differentiation protocol. Doses of 0.75 µM retinoic acid (RA) were added to the cells every 24 hours on days 4, 5, and 6 with media exchanged to RPMI1640 + B27 with insulin at day 7. Cells were harvested for analysis at day 20. B) qPCR analysis of ventricular markers *MYL2* and IRX4, cardiac marker *NKX2*.*5*, and atrial markers *NPPA, GJA5, CACNA1D, KCNA5*, and *KCNJ3*. n = 3, unpaired t-test, *p < 0.05. C) Flow cytometric analysis of cardiac troponin T (cTnT) and myosin light chain 2v (normalized to cTnT expression) in hiPSC-aCMs and -vCMs. n = 4, unpaired t-test, ***p<0.001. D) Average beating rates of hiPSC-aCMs and -vCMs from the day they begin to beat until day 20. n = 4 independent differentiation batches E) Atrial Natriuretic peptide (ANP) concentration between hiPSCs, and hiPSC-aCMs and -vCMs determined by competitive ELISA. n = 3 and n = 2 hiPSC lines, unpaired t-test *p<0.05, ** p<0.01, ***p<0.001. Data are presented as mean ± SEM and the n represents the number of independent differentiation batches.

Compared to hiPSC-vCMs, hiPSC-aCMs were found to have no significant difference in the pan cardiac phenotype. Expression of the pan cardiac transcript *NKX 2*.*5* measured by qPCR was similar between hiPSC-aCMs and -vCMs (Figure 1B), as was cardiac troponin T (cTnT) protein expression measured by flow cytometry (Figures 1C and S1). The protein expression of MLC-2v was reduced in hiPSC-aCMs compared to hiPSC-vCMs (8.0 ± 1.1 % vs. 57.0 ± 0.5 %; p<0.05) (Figure 1C). Furthermore, hiPSC-aCMs displayed higher concentrations (increased by 91%) of atrial natriuretic peptide (ANP) at 65 ± 2 compared to 34 ± 6 ng/mL in hiPSC-vCMs as measured by ELISA (p<0.05).

The qPCR assay revealed that atrial-specific transcripts such as atrial natriuretic peptide (*NPPA*), connexin 40 (*GJA5*), the L-type calcium channel CaV1.3 (*CACNA1D*), and the K^+^ channels K_v_1.5 (*KCNA5*) and K_ir_3.1 (*KCNJ3)* transcripts were all expressed at a significantly higher levels in hiPSC-aCMs compared to hiPSC-vCMs (p<0.05, Figure 1B). Another ventricular marker, *IRX4*, also had decreased expression in hiPSC-aCMs (Figure 1B). Furthermore, consistent with previous studies^8–10,14,15^, hiPSC-aCMs started beating at day 10 or earlier and exhibited an increased beating frequency relative to hiPSC-vCMs, which started beating around day 10-12 post-differentiation.

### Gene Expression Analysis of hiPSC-aCMs

We performed an extensive gene expression analysis of hiPSC-aCMs and -vCMs using NanoString technology in which each mRNA copy was digitally counted for accurate and sensitive detection of gene expression^16^. Five independent differentiation batches of each cardiac subtype were included in the analysis. The unsupervised hierarchical clustering analysis showed clear grouping of hiPSC-aCM samples that were segregated relative to hiPSC-vCMs (Figure 2A). The gene expression profile of the hiPSC-vCM samples were more variable with 2 samples closer in distance to the hiPSC-aCMs while 3 samples displayed clear segregation (Figure 2A). The overall difference in global gene expression and lineage between hiPSC-aCMs and -vCMs was also captured in the principal component analysis (PCA, Figure S4A). Out of the 250 transcripts analyzed, 200 genes were detected above background noise defined by a threshold of 50 raw digital counts as determined by the negative controls of the assay. In the hiPSC-aCMs, 14 and 27 genes were significantly upregulated and downregulated, respectively (Figure 2C). As expected, hiPSC-aCMs displayed significantly higher expression profiles of atrial-specific markers including atrial-specific K^+^ channel Kv1.5 (*KCNA5*) and transcription factors (*NR2F2* and *TBX18*) (Figure 2C). Meanwhile, hiPSC-vCMs displayed higher expression of ventricular-specific genes such as those encoding for contractile proteins *MYL2, MYH7*, and the L-type Ca2+ channel isoform Ca_v_1.2 (*CACNA1C*) (Figure 2C). The genes encoding for the proteins in the sarcoplasmic reticulum complex such as *TRDN, CASQ2*, and *RYR2* were expressed in significantly lower amounts in the hiPSC-aCMs samples (Figure 2C). Meanwhile, pan-cardiac markers *NKX2-5* and *TNNT2* were expressed at similar levels in both hiPSC-aCMs and -vCMs, further corroborating the efficacy of the differentiation protocol (Figure S4B).

**Figure 2:**
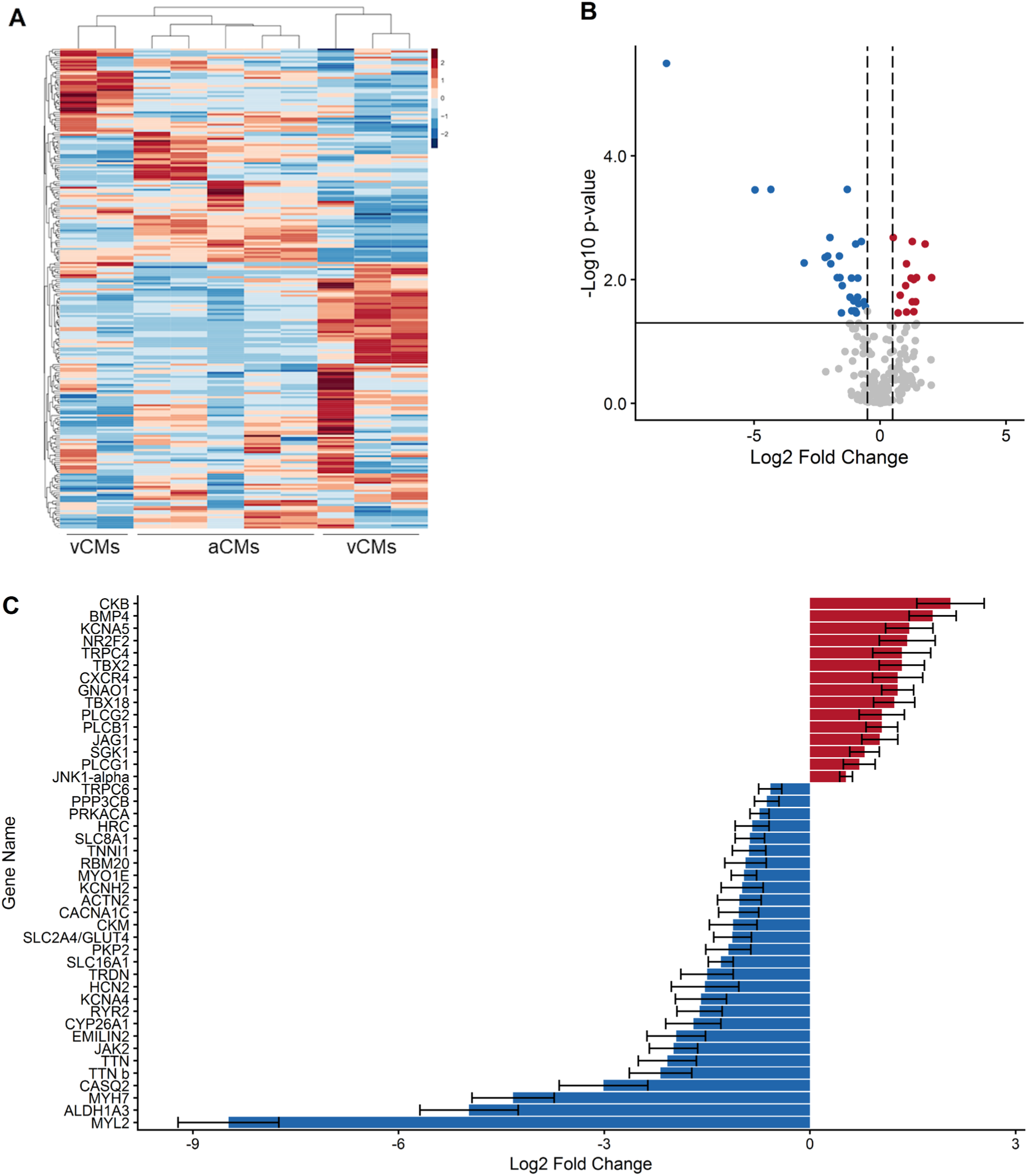
Gene expression analysis of hiPSC-aCMs and -vCMs using NanoString. Global gene expression pattern of hiPSC-aCMs and -vCMs shown in A) heat map of the expression of the 250 genes across samples of hiPSC-aCMs and -vCMs. The cluster dendrogram shows the unsupervised hierarchical clustering that was conducted using the agglomerative algorithm and the Euclidian distance criterion. B) Differentially expressed genes between hiPSC-aCMs and -vCMs expressed in volcano plot shows 14 upregulated (red) and 27 downregulated (blue) genes in hiPSC-aCMs. Solid horizontal line represents the Benjamini-Hochberg false discovery rate (FDR) adjusted p-value < 0.05 (-log10 = 1.3). Dashed vertical lines represent the arbitrary log2 fold change cut-off of -0.5 and 0.5. C) 42 differentially expressed genes identified from the statistical criteria of FDR adjusted p-value < 0.05 and log2 fold change of <-0.5 and > 0.5. Data are presented as mean ± SEM. n = 5 independent differentiation batches.

**Figure 3:**
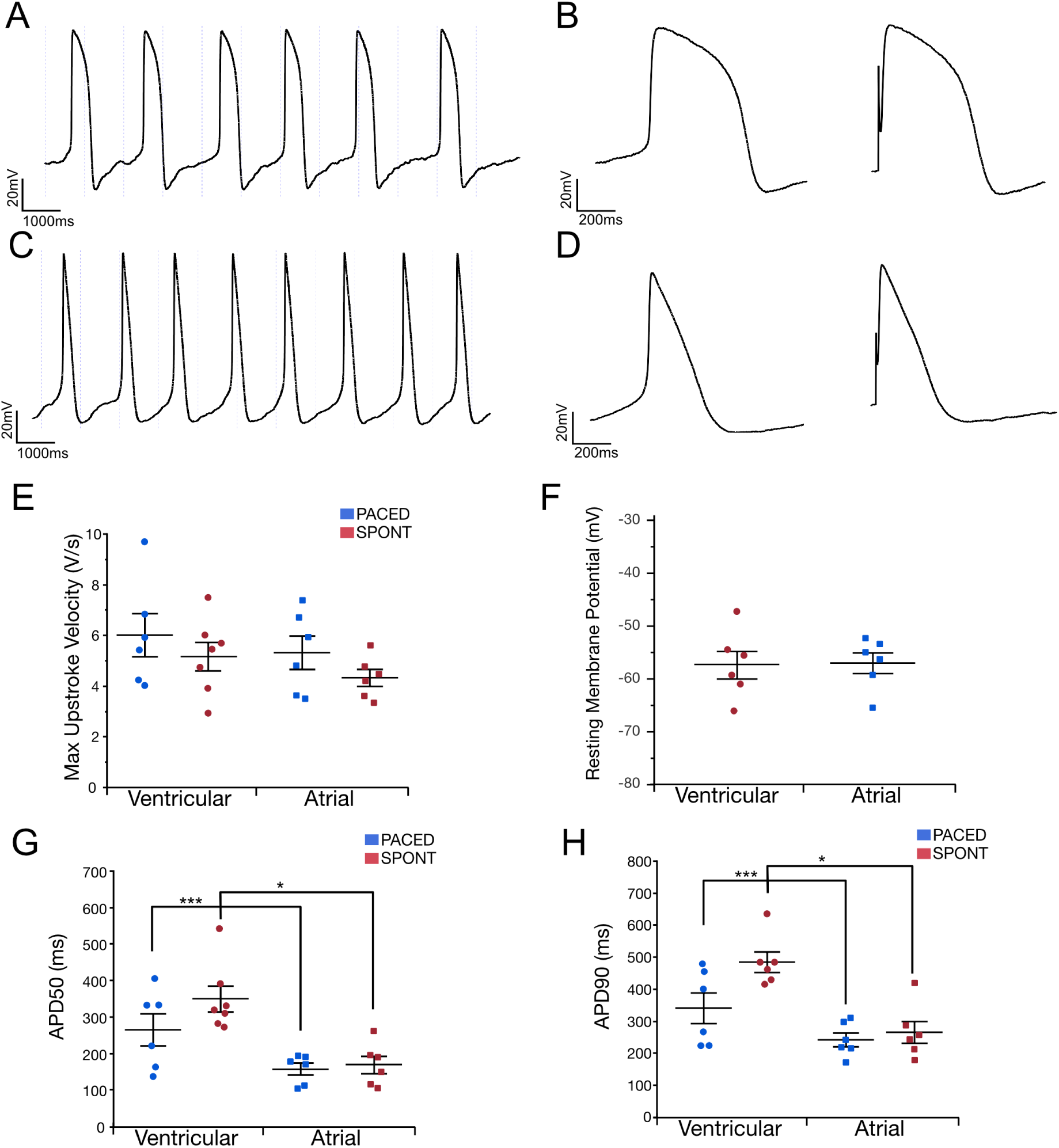
hiPSC-aCMs and -vCMs have distinct electrophysiological characteristics. Single differentiated hiPSC-aCMs and -vCMs were plated on gelatin and Geltrex after 30 days in culture. A) Whole cell current clamp recordings from a spontaneously beating hiPSC-vCM. B) Recorded action potential (APs) demonstrates typical prolonged plateau phase in both spontaneous (left) and/or paced at 1 Hz (right). C) Current clamp recording from a spontaneously beating hiPSC-aCM. D) Single AP from hiPSC-aCM demonstrates shortened action potential duration (APD) and lack of prolonged plateau phase, spontaneous (left), paced at 1 Hz (right). E) The first differential of voltage recordings from hiPSC-aCMs and –vCMs were used to calculate the maximal upstroke velocities. F) One minute after achieving the whole-cell configuration, the average resting membrane potential was measured. G) Spontanous and 1 Hz paced APs were assessed for duration at 50% of peak (APD_50_), and H) 90% of peak (APD_90_). Statistics were performed by unpaired t-test. * p < 0.05, *** p <0.005. Data are presented as mean ± SEM. Two differentiation batches were included in this analysis.

### Functional Phenotyping of hiPSC-derived Atrial Cardiomyocytes

We compared the electrophysiological characteristics of the differentiated hiPSC-aCMs and –vCMs using whole-cell patch clamp. Confirming our observations in tissue culture, the spontaneous beating rates were higher in the single hiPSC-aCMs than in –vCMs (Figure A and C). Whole cell current clamp recordings demonstrated the ventricular-like action potential (AP) morphology of hiPSC-vCMs with a clear and prolonged plateau phase while the AP of the hiPSC-aCMs displayed atrial-like morphology with a shorter action potential duration (APD) and a lack of prolonged plateau phase at both spontaneous beating rates (Figure B & D, left panel) and paced at 1 Hz (Figure B & D, right panel). No statistical differences were observed in the resting membrane potential and the maximum upstroke velocity of hiPSC-aCMs and –vCMs. The APD at 50% (APD_50_) and 90% (APD_90_) of the peak voltage were significantly shorter in hiPSC-aCMs than –vCMs at both spontaneous beating rates (APD_50_: 157 ± 16 ms vs. 349 ± 35 ms, p-value < 0.005; APD_90_: 249 ± 34 ms vs. 484 ± 30 ms, p-value < 0.005) and paced at 1 Hz (APD_50_: 157 ± 16 ms vs. 264 ± 44 ms, p-value < 0.05; APD_90_: 242 ± 22 ms vs. 341 ± 48 ms, p-value < 0.05).

We further assessed the functional properties of hiPSC-aCMs and –vCMs using optical mapping with simultaneous measurement of APs and calcium transients (CaT). Like the patch clamp recordings, optical membrane voltage measurements revealed similar atrial-like and ventricular-like AP morphology in the hiPSC-aCMs and –vCMs, respectively (Figure 4A). AP and CaT durations were quantified at early, mid, and late repolarization (APD_20_, APD_50_, and APD_80_) and Ca2+ decay (CaTD_20_, CaTD_50_, and CaTD_80_), respectively. These stages reflect different phases of ionic currents across the plasma membrane and the extrusion of Ca^2+^ handling mechanics.

**Figure 4:**
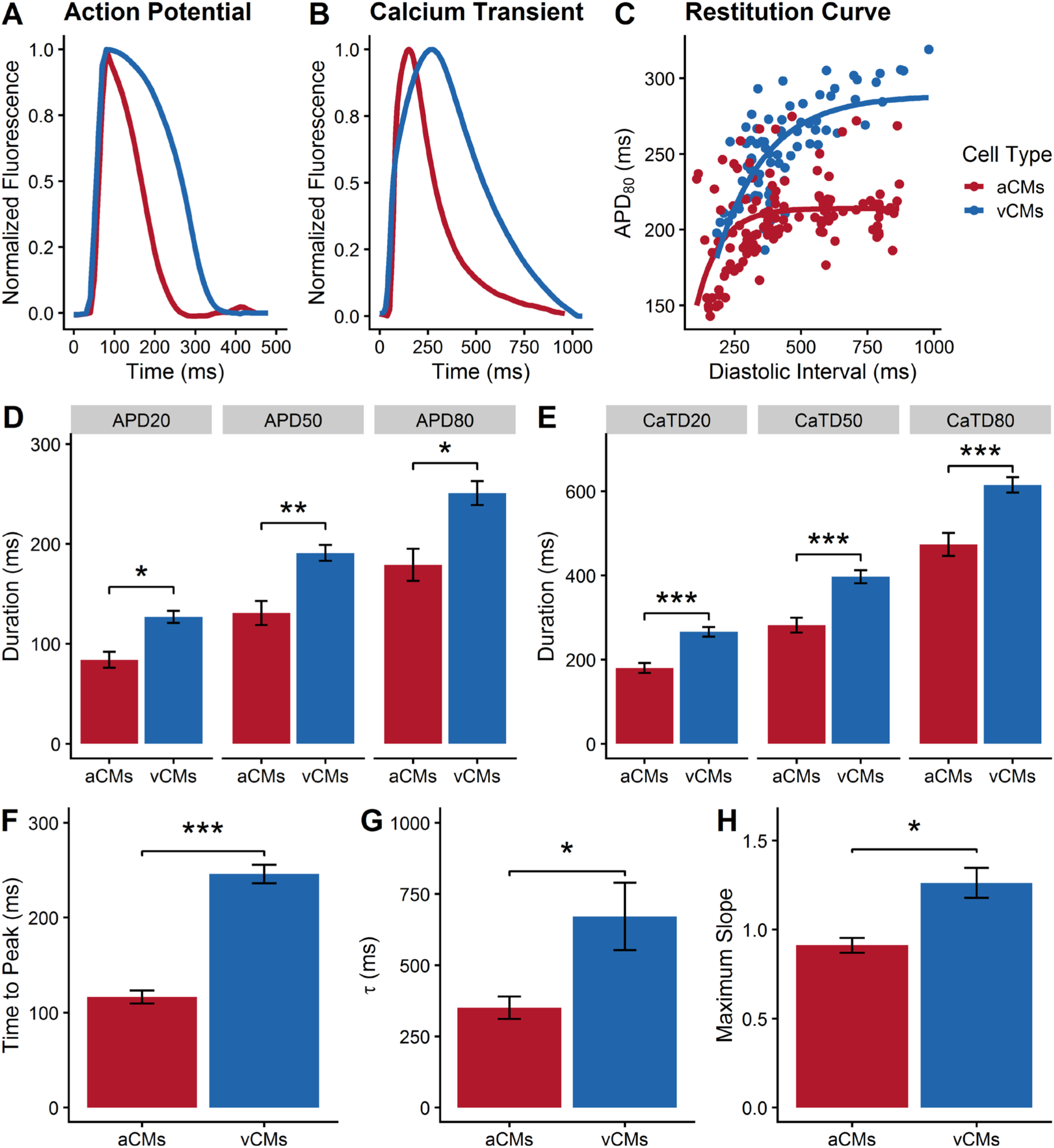
Functional phenotyping of hiPSC-derived atrial and ventricular CMs using optical mapping. Representative average traces of A) action potential and B) Ca^2+^ transients of hiPSC-aCMs and –vCMs electrically paced at 1 Hz. C) Electrical restitution curve measured at APD_80_ relative to the diastolic interval (DI). D) Quantification of early- (APD_20_), mid- (APD_50_), and late- (APD_80_) repolarization, unpaired t-test, *p < 0.05, **p < 0.01. E) Quantification of early- (CaTD_20_), mid- (CaTD_50_), and late- (CaTD_80_) Ca2+ transient decay, unpaired t-test, ***p < 0.001. F) Time to peak (TTP) of the Ca^2+^ transient, unpaired t-test, ***p < 0.001. G) Time constant (τ) of Ca^2+^ decay, unpaired t-test *p < 0.05. H) Maximum slope of the electrical restitution as shown in panel C, unpaired t-test, *p < 0.05. Electrical restitution curves were measured under a variable rate pacing protocol (60 – 200 bpm) as described in the Supplemental information. n = 4 (four independent differentiation batches) and cardiac enriched hiPSC-aCMs and -vCMs were analyzed in these set of experiments. Data are presented as mean ± SEM.

For these experiments, both hiPSC-aCMs and -vCMs were paced at 1 Hz. All measured levels of the APD were significantly shorter in hiPSC-aCMs compared to hiPSC-vCMs (APD_20_: 84 ± 8 ms vs. 127 ± 6 ms, p < 0.05; APD_50_: 131 ± 12 ms vs. 191 ± 8 ms, p < 0.01; APD_80_: 179 ± 16 ms vs. 251 ± 12 ms, p < 0.05; Figure 4D). The overall CaTD of hiPSC-aCMs was significantly shorter than that of hiPSC-vCMs (CaTD_20_: 180 ± 12 ms vs. 266 ± 12 ms, p < 0.001; CaTD_50_: 282 ± 18 ms vs. 397 ± 16 ms, p < 0.001; CaTD_80_: 474 ± 27 ms vs. 615 ± 18 ms, p < 0.001; Figure 4E). Compared to hiPSC-vCMs, hiPSC-aCMs displayed significantly faster CaT time-to-peak (hiPSC-aCMs: 116 ± 7 ms vs. hiPSC-vCMs: 246 ± 10 ms, p < 0.05) and faster decay kinetics (τ; hiPSC-aCMs: 350 ± 39 ms vs. hiPSC-vCMs: 671 ± 118 ms, p < 0.05) indicating that Ca^2+^ handling mechanics are accelerated in hiPSC-aCMs (Figure 4F & G).

The direct comparison between whole-cell patch clamp and optical mapping read-outs paced at 1 Hz is shown in Figure S7. We observed no differences in the read-outs of hiPSC-aCMs at APD_20_ (optical: 84 ± 8 ms, patch: 98 ± 12 ms) and APD_50_ (optical: 131 ± 12 ms, patch: 169 ± 19 ms). However, APD_80_ of hiPSC-aCMs measured by patch clamp was longer than the optical APD_80_ (253 ± 22 ms vs. 179 ± 16 ms, p < 0.05). Similarly, both APD_20_ (216 ± 22 ms vs. 127 ± 6 ms) and APD_80_ (393 ± 62 vs. 251 ± 12 ms) of hiPSC-vCMs measured by patch clamp were longer than the comparable optical measurements. APD_50_ of hiPSC-vCMs did not show a statistical difference between the two recording paradigms (optical: 191 ± 8 ms, patch: 308 ± 60 ms).

Rate-dependent properties are critical in cardiac function. A variable rate protocol (Figure S6) in which the hiPSC-CMs were electrically paced with increasing frequency at every cycle was used to investigate the electrical restitution dynamics. The electrical restitution curve reflects the ability of the cardiac system to accommodate a higher pacing rate by progressive shortening of APD_80_ and is described as APD_80_ in relation to the diastolic interval (DI).

Compared to hiPSC-vCMs, the electrical restitution curve of the hiPSC-aCMs displayed a flatter portion and did not show APD_80_ shortening at longer diastolic intervals (Figure 4F). The extensive shortening in APD_80_ started at shorter diastolic intervals for hiPSC-aCMs (< 275 ms) compared to hiPSC-vCMs (<500 ms). The maximum slope of the restitution curve was higher in hiPSC-vCMs compared to hiPSC-aCMs (1.26 ± 0.08 vs. 0.91 ± 0.04, p < 0.05; Figure 4G) indicating faster kinetics of APD in response to higher pacing rate.

### *In Vitro* Screening for Atrial-selective Pharmacology

We first established the utility of optical mapping to detect a pan-cardiac pharmacological response by using dofetilide, a strong blocker of the rapid delayed rectifier K^+^ current (I_Kr_)^17^, an ionic current expected to be present in both hiPSC-aCMs and –vCMs^18^. Dofetilide elicited a dose-dependent response in both hiPSC-aCMs and –vCMs. Compared to pre-drug baseline, dofetilide at 100 nM prolonged APD_80_ of both hiPSC-aCMs from 182 ± 16 ms to 355 ± 24 ms (95 + 7 % prolongation) and of hiPSC-vCMs from 238 ± 20 ms to 319 ± 45 ms (34 ± 14 % prolongation, p < 0.05; Table S1 & Figure 5C). The drug prolonged early-repolarization (APD_20_) of hiPSC-vCMs at 10 and 30 nM while having no effect on APD_20_ of hiPSC-aCMs at all tested doses (Table S1). Additionally, CaTD_50_ and CaTD_80_ of both hiPSC-aCMs and –vCMs were significantly prolonged in response to dofetilide (Table S1). However, hiPSC-aCMs appeared to be more sensitive to dofetilide as the APD_80_ was significantly prolonged at the lowest tested dose of 3 nM (from to 182 ± 26 ms to 241 ± 26 ms, p < 0.05; Table S1) and displayed a larger dose-response (Figure S8).

**Figure 5:**
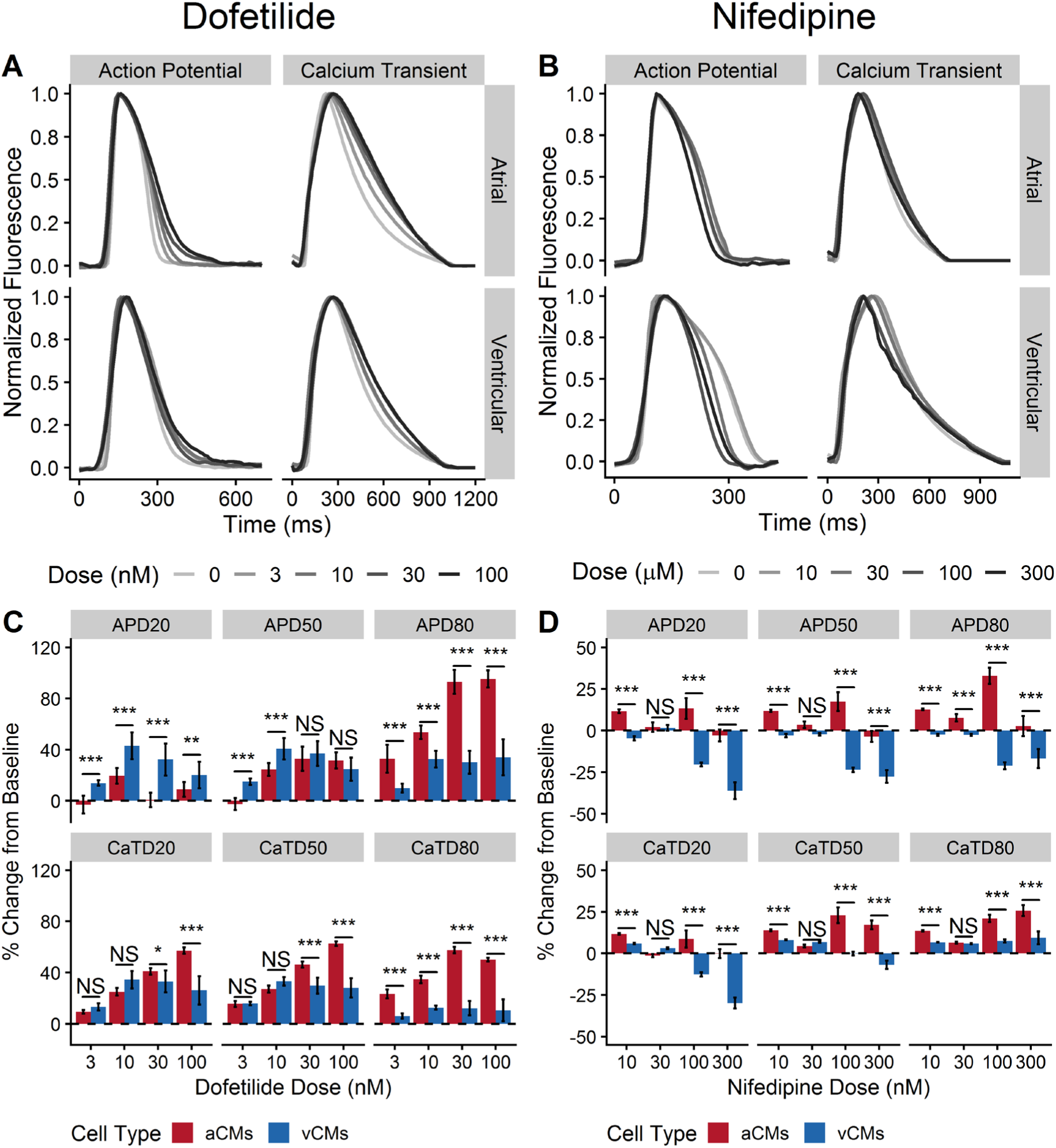
The effects of dofetilide and nifedipine on action potential and Ca^2+^ transient of hiPSC-aCMs and -vCMs. Representative traces of action potential and Ca^2+^ transients illustrating the effects of A) dofetilide and B) nifedipine on hiPSC-aCMs and –vCMs. Higher drug doses are presented by a progressively darker shade. The effects of C) 4-aminopyridine and D) nifedipine on normalized (percent change from pre-drug baseline) action potential duration (APD) and Ca^2+^ transient duration (CaTD); both parameters being measured at 20, 50, and 80%. Dashed line is the normalized pre-drug control presented as 0% change. n = 6 from six independent differentiation batches. hiPSC-derived atrial cardiomyocytes (aCMs) are shown in red while hiPSC-derived ventricular cardiomyocytes (vCMs) are presented in blue. Data are presented as mean ± SEM. Drug effects were compared between hiPSC-aCMs and -vCMs at each dose using unpaired t-test, *p < 0.05, **p < 0.001, ***p < 0.001. NS stands for not significant.

Next, we demonstrated the functional differences in the ion channels of hiPSC-aCMs and -vCMs. We aimed to show that the ultra-rapid outward current (I_Kur_) produced by the channel K_v_1.5 (*KCNA5*) was functional and specific to hiPSC-aCMs, while the inward Ca^2+^ current (I_Ca,L_) produced the voltage-dependent L-type Ca^2+^ channel Ca_V_1.2 (*CACNA1C*) was functional and specific to hiPSC-vCMs. We used two relatively selective compounds, 4-aminopyridine (4AP) and nifedipine, to dissect the presence of functional I_Kur_ and I_CaL_, respectively. While nifedipine is also known to block Cav1.3, it has been to have a preferential effect at lower concentrations on Ca_V_1.2 with upwards of ∼13-fold higher block on Ca_V_1.2 than Ca_V_1.3^19^.

At the highest tested dose (300 nM), nifedipine significantly decreased APD_50_ of hiPSC-vCMs from 170 ± 14 ms to 121 ± 16 ms (28% shortening) and decreased CaTD_50_ from 357 ± 10 ms to 333 ± 23 ms (30% shortening) (Figure 5D; Table S2). We observed a trend in APD_50_ shortening of hiPSC-aCMs in response to increasing nifedipine doses, but the drug elicited a significantly stronger dose-dependent shortening in both APD and CaTD of hiPSC-vCMs compared to hiPSC-aCMs (Figure S8 & S9). Observing the percent change from pre-drug control, nifedipine induced differential response in overall APD and CaTD between hiPSC-aCMs and –vCMs at 10, 100, and 300 nM (Figure 5D).

In hiPSC-aCMs, 4AP prolonged APD and CaTD in a dose-dependent manner with a statistically significant change starting at 30 µM (Figure 6A & C; Table S3). 4AP significantly prolonged early-repolarization (APD_20_) of hiPSC-aCMs by 46 ± 2 % and 66 ± 2 % at 50 and 100 µM, respectively (APD_20_ at baseline: 82 ± 8, at 50 µM: 120 ± 9 ms, at 100 µM: 131 ± 9 ms, p < 0.05) (Figure 6C & Table S3). In contrast, 4AP prolonged APD_20_ of hiPSC-vCMs by 23% (APD_20_ at baseline: 138 ± 8 ms, at 100 µM: 170 ± 9 ms) at the highest tested dose of 100 µM (Figure 6C & Table S3). hiPSC-aCMs showed greater change in APD to relative to pre-drug control at all concentrations of 4AP compared to hiPSC-vCMs (Table S3), This is corroborated by the steeper trend of the dose response relationship in hiPSC-aCMs (Figure S8). Additionally, the overall CaTD of hiPSC-aCMs were prolonged after exposure to 4AP at 10 µM while the drug had a significant effect on CaTD of hiPSC-vCMs at 30 µM (CaTD_50_ elongation from baseline: 68 ± 2 % vs. 12 ± 2 %, p < 0.05) (Table S3).

**Figure 6:**
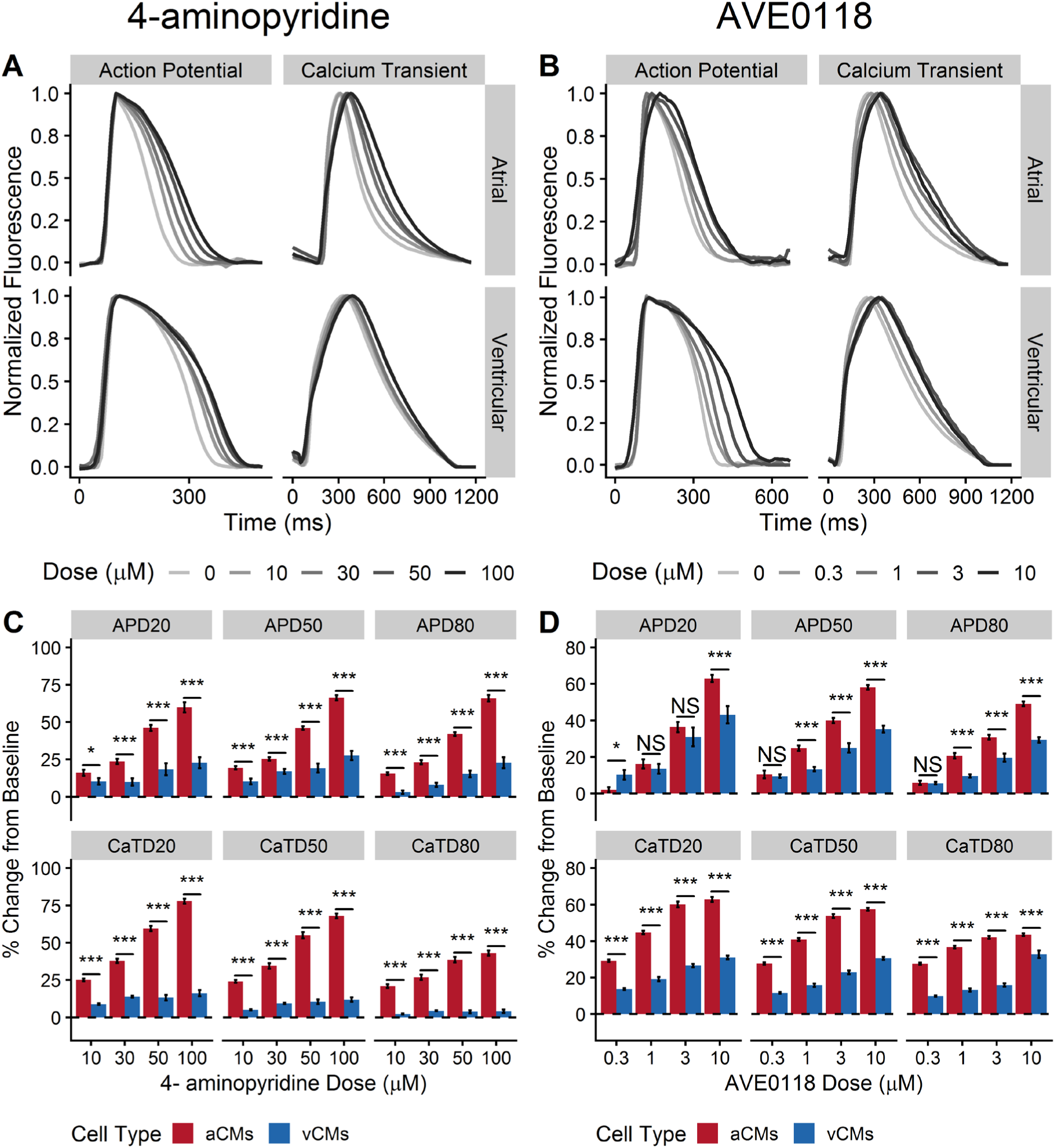
The effects of 4-aminopyridine (4AP) and AVE0118 on action potential and Ca^2+^ transient of hiPSC-aCMs and -vCMs. Representative traces of action potential and Ca^2+^ transients illustrating the effects of A) 4-aminopyridine (4AP) and B) AVE0118 on hiPSC-aCMs and –vCMs. Higher drug dose is presented by a progressively darker shade. The effects of C) dofetilide and D) vernakalant on normalized (percent change from pre-drug baseline) action potential duration (APD), and B) Ca^2+^ transient duration (CaTD); both parameters being measured at 20, 50, and 80%. Dashed line is the normalized pre-drug control presented as 0% change. n = 6 from six independent differentiation batches. hiPSC-derived atrial cardiomyocytes (aCMs) are shown in red while hiPSC-derived ventricular cardiomyocytes (vCMs) are presented in blue. Data are presented as mean ± SEM. Drug effects were compared between hiPSC-aCMs and -vCMs at each dose using unpaired t-test, *p < 0.05, **p < 0.001, ***p < 0.001. NS stands for not significant.

We then demonstrated the effectiveness of our drug screening platform in assessing the effects of experimental compounds designed to have targeted effects on atrial-specific ion channels using AVE0118 and UCL1684.

AVE0118 is an experimental drug that blocks I_Kur_, the G-protein-activated K^+^ current (I_KAch_), and the transient outward K^+^ current (I_to_) at a similar dose range^20^. Both I_Kur_ and I_KAch_ are atrial-specific ionic currents. AVE0118 prolonged mid- and late-repolarization (APD_50_ and APD_80_) of both hiPSC-aCMs and –vCMs at the two highest tested doses (3 and 10 µM; Table S4). Similarly, AVE0118 had significant effects on CaTD_50_ and CaTD_80_ of hiPSC-aCMs and – vCMs at all tested doses (Table S4). However, the APD_50_ and APD_80_ of hiPSC-aCMs were significantly prolonged at a lower dose of 1 µM (control: 200 ± 14 ms, 1 µM: 244 ± 16 ms; Table S4). Furthermore, the atrial-selective effects of the drug were demonstrated by a larger proportional prolongation in APD_50_ and APD_80_ of hiPSC-aCMs compared to hiPSC-vCMs at 1, 3, and 10 µM (APD; Figure 6D). Furthermore, AVE0118 induced a larger proportional prolongation in CaTD of hiPSC-aCMs compared to hiPSC-vCMs at all tested doses (Figure 6D). Early repolarization (APD_20_) of hiPSC-aCMs also displayed a large dose-dependent response (Figure S8) with a proportionally larger prolongation at 10 µM (63 ± 2% vs. 43 ± 5%, p < 0.05; Figure 6D)

UCL1684 is purported to be a potent direct pore blocker of the small conductance Ca^2+^ activated K^+^ channel (SK channel)^21^ and was expected to induce a dose-dependent atrial-selective response. In hiPSC-aCMs, UCL1684 treatment resulted in a significantly prolonged APD_80_ at 3 µM and 10 µM (from pre-drug control: 136 ± 11 ms to 3 µM: 188 ± 25 ms or 38% prolongation, and to 10 µM: 206 ± 32 ms or 49% prolongation, p < 0.05; Figure 7C & Table S5). UCL1684 prolonged CaTD_80_ of hiPSC-aCMs at all tested doses (baseline: 300 ± 15 ms, at 0.3 µM: 372 ± 23 ms, at 1 µM: 387 ± 33 ms, at 3 µM: 413 ± 24 ms, at 10 µM: 416 ± 39 ms, p < 0.05; Table S5). In contrast, UCL1684 exposure showed no statistically significant effect on overall APD and CaTD of hiPSC-vCMs. The sensitivity of hiPSC-aCMs to UCL1684 was also reflected in the dose-response relationship showing a prolongation APD_80_, in contrast to the minimal prolongation in APD_80_ of hiPSC-vCMs (Figure S8).

**Figure 7:**
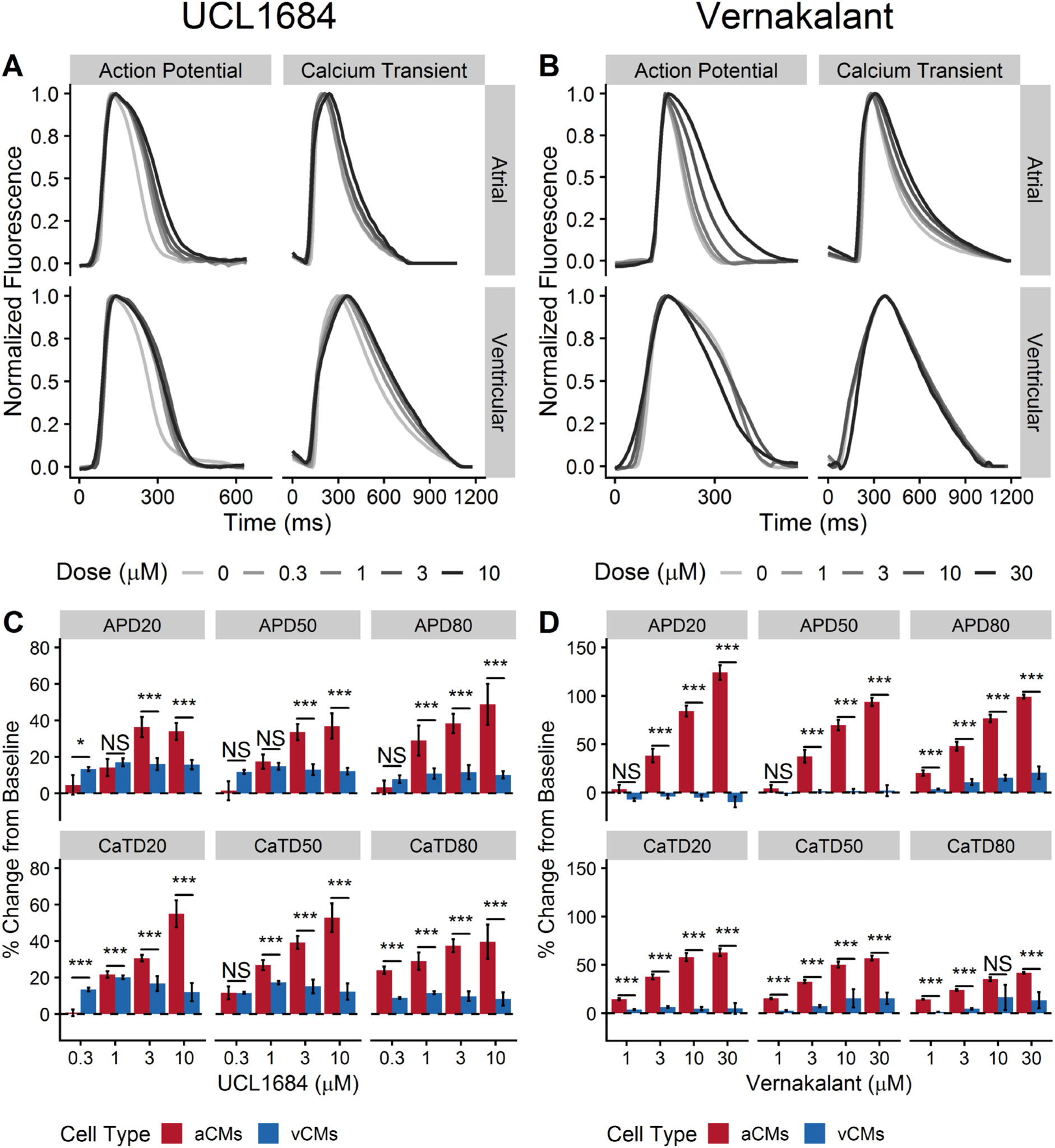
The effects of UCL1684 and vernakalant on action potential and Ca^2+^ transient of hiPSC-aCMs and -vCMs. Representative V_m_ and Ca^2+^ transients illustrating the effects of A) UCL1684 and B) vernakalant on hiPSC-aCMs and –vCMs. Higher drug doses are presented by a progressively darker shade. The effects of C) AVE0118 and D) UCL1684 on normalized (percent change from pre-drug baseline) action potential duration (APD) and Ca^2+^ transient duration (CaTD); both parameters being measured at 20, 50, and 80%. Dashed line is the normalized pre-drug control presented as 0% change. n = 6 from six independent differentiation batches. hiPSC-derived atrial cardiomyocytes (aCMs) are shown in red while hiPSC-derived ventricular cardiomyocytes (vCMs) are presented in blue. Data are presented as mean ± SEM. Drug effects were compared between hiPSC-aCMs and -vCMs using unpaired t-test at each dose, *p < 0.05, **p < 0.001, ***p < 0.001. NS stands for not significant.

Finally, we tested the effects of vernakalant which is a multi-ion channel blocker that blocks the fast and late inward Na^+^ current (I_Na_, I_NaL_, respectively), the I_Kur,_ and the I_KAch_ ^22^. The drug is used clinically for intravenous cardioversion of patients in AF ^23^ and was expected to induce an atrial-specific effect due to its I_Kur_ and I_KAch_ blocking properties.

Vernakalant elicited a positive dose-dependent response in both APD and CaTD of hiPSC-aCMs with minimal measurable effects on hiPSC-vCMs (Table S6; Figure S8 & S9). Vernakalant demonstrated atrial-selectivity with statistically significant differences between APD and CaTD of hiPSC-aCMs and –vCMs at doses of 3, 10, and 30 µM (Figure 7D). Compared to APD at baseline, vernakalant at 10 µM significantly prolonged APD_20_, APD_50_, and APD_80_ of hiPSC-aCMs by 84%, 70 %, and 77%, respectively (Figure 7D). Additionally, vernakalant at 10 µM prolonged CaTD_20_, CaTD_50_, CaTD_80_ of hiPSC-aCMs by 58%, 50%, 35%, respectively (Figure 7D). At clinically relevant concentrations (30 µM), vernakalant greatly affected early repolarization of hiPSC-aCMs (APD_20_ prolonged by 124%; Figure 7D). At 30 µM, vernakalant prolonged APD_80_ of hiPSC-vCM by 20% (APD_80_: 238 ± 22 ms at baseline vs. 289 ± 30 ms at 30 µM, p < 0.05; Figure 7D & Table S6). Except for APD_80_ prolongation at 30 µM, vernakalant had no statistically significant effect on overall APD and CaTD of hiPSC-vCMs at the lower doses (Table S6).

## Discussion

In this study, we were successful in efficiently differentiating hiPSCs into a monolayer of cardiomyocytes with an atrial phenotype by modifying the GiWi protocol^11^. We used multiple phenotypic approaches such as qPCR, digital multiplexed gene expression analysis with NanoString technology, flow cytometry, ELISA, voltage measurements with current clamp electrophysiology as well as simultaneous voltage and Ca^2+^ transient measurements with optical mapping to demonstrate a clear and distinct atrial phenotype. Unique to our study, we completed an in-depth pharmacological analysis with simultaneous voltage and Ca^2+^ measurements to demonstrate the differential responses of these chamber-specific cardiomyocytes, and their utility as a translational model in screening for the safety and efficacy of novel atrial-specific compounds for the treatment of AF.

Our observations support previous data in showing that atrial specification is in part mediated by RA^6,8–10,15^. In our protocol, atrial differentiation was accomplished by adding 0.75 µM RA twenty-four hours after WNT inhibition, with a total exposure time of 72 hours. The generated hiPSC-aCMs showed an atrial-specific phenotype as validated at both protein and transcript levels with a decrease in ventricular-specific and an increase in atrial-specific markers. These results suggest that RA, at the dose and temporal exposure used in this study, maintains cardiac differentiation efficacy while pushing the differentiation process into an atrial lineage.

As a complementary assay, we used the NanoString digital multiplexed gene expression analysis to assess the expression of 250 genes custom-curated from the existing literature. We found *MYL2* and *MYH7*, markers of the ventricular phenotype, to be significantly differentially expressed between hiPSC-aCMs and –vCMs, matching the gene expression pattern of native adult human right atrial and left ventricular tissues^24^. Another ventricular-specific marker *KCNA4*^25^ which encodes for the Kv1.4 channel of the slow I_to_ was downregulated in hiPSC-aCMs. Canonical atrial markers such as *KCNA5* and *NR2F2* were also confirmed to be differentially upregulated in hiPSC-aCMs. Other markers of human atrial specificity such *CXCR4, GNAO1, JAG1, PLCB1, and TBX18* as retrieved from the GTEx database^26^ were upregulated in our hiPSC-aCMs further demonstrating the effect of RA on driving the differentiation pathway into an atrial lineage.

*MYL7*, thought to be an atrial-specific marker, was not found to have a significantly higher expression in hiPSC-aCMs. The differential expression of MLC-2a may however require additional maturation of the hiPSC-CMs. Other studies^11,27^ have shown a high expression in MLC-2a at day 20 post-differentiation and a subsequent decrease over time in culture systems generating predominantly ventricular hiPSC-CMs. One study has shown a higher expression of MLC-2a in hiPSC-aCMs analyzed at a later date (earliest at day 60)^6^.

Electrophysiological differences between atrial and ventricular cardiomyocytes, in terms of voltage and Ca^2+^ handling, define their function and are critical to the development and determination of efficacy of atrial-specific compounds. As demonstrated by whole-cell patch clamp and optical mapping measurements, the hiPSC-aCMs generated in this study exhibited atrial-like AP and Ca^2+^ handling properties. Namely, the AP of hiPSC-aCMs were significantly shorter, along with a lack of a prolonged plateau phase as opposed to the AP of hiPSC-vCMs, an observation that is aligned with native cardiomyocyte electrophysiology^28^. Similarly, the CaT of hiPSC-aCMs had faster kinetics with a faster decay time as reflected by the differential expression of Ca^2+^ channel isoforms, further demonstrating the differential physiology between hiPSC-aCMs and -vCMs.

In terms of APD measurements, we observed a good correlation between the patch clamp and optical mapping recordings for hiPSC-aCMs. In hiPSC-vCMs, however, the optical AP measurements were shorter overall than patch clamp recordings. This discrepancy may be attributed to the heterogeneity of our current ventricular differentiation protocol which generated predominantly ventricular cardiomyocytes but also contain a small proportion of non-ventricular phenotypes (i.e. atrial myocytes and nodal cells). Thus, the optical AP signals in the rig used represent an average from about 300,000 cells in each 1 cm^2^ region of interest.

Another hallmark of cardiomyocyte function is rate-dependence, as described by the electrical restitution curve^29^. We observed that the electrical restitution properties were different between hiPSC-aCMs and -vCMs. Compared to hiPSC-vCMs, hiPSC-aCMs displayed a steady-state-like property by undergoing minimal APD_80_ shortening in response to the lower ranges of the pacing protocol (cycle lengths of about 400 to 1000 ms) indicating full recovery of ion channel kinetics at these pacing ranges. In contrast, the hiPSC-vCMs displayed consistent APD_80_ shortening at the same pacing range. It is important to note that APD restitution curves are likely different when using the standard steady-state extra stimulus protocol compared to dynamic pacing, particularly in cardiomyocytes with immature Ca^2+^ handling and memory^29^. In relation to dynamic pacing protocol, hiPSC-vCMs exhibit a steeper maximum slope of the restitution curve compared to hiPSC-aCMs as steady-state APD is the principal determinant of the slope of the ventricular restitution curve^30^.

The presence of specific ion channel currents (i.e. I_Kur_, I_KAch_, and I_CaL_) explain, in part, the functional differences between the two cardiac chamber sub-types, the expressions of which were already shown in our qPCR and NanoString assays. We used a series of compounds (4-aminopyridine, dofetilide, vernakalant, AVE0118, UCL1684, and nifedipine) to demonstrate the function of atrial-specific ionic currents in our model system and were able to show the expected chamber specific differences between hiPSC-aCMs and -vCMs.

Dofetilide (DF) served as a positive control in our optical mapping assay as a clinically relevant drug which has a strong effect on I_Kr_ in both atria and ventricular CM^31^. As expected, dofetilide affected the repolarization of both hiPSC-aCMs and –vCMs, confirming the presence of I_Kr_ in both cell types. At clinically relevant doses of DF (3 and 10 nM), hiPSC-aCMs displayed greater sensitivity to the drug indicating a larger proportional contribution of I_Kr_ in the AP of hiPSC-aCMs relative to hiPSC-vCMs. This may partly explain the effectiveness of the drug in the clinical treatment of AF. However, clinical use of the drug to treat AF is limited due to its tendency to induce QT_c_ prolongation. This pro-arrhythmic risk of TdP^32^ which was captured by the prolongation of APD_80_, an *in vitro* surrogate of QT_c_, in the hiPSC-vCMs. This finding supports the utility of our optical mapping assay in predicting the risk of ventricular arrhythmogenesis *in vitro*.

The compound 4AP has been shown to selectively block K_v_1.4 (I_to_) and K_v_1.5 (I_Kur_)^33^ and is therefore expected to elicit a response in hiPSC-aCMs at lower doses than in hiPSC-vCMs as I_Kur_ (K_v_1.5) is a strong functional indicator of atrial phenotype. Confirmation of the atrial expression of I_Kur_ channels was demonstrated by the stronger dose-dependent hiPSC-aCM AP prolongation to 4AP at all tested doses (10, 30, 50 and, 100 µM) suggesting selective sensitivity of hiPSC-aCMs to 4AP due to a greater expression of K_v_1.5. The inhibitory effects of 4AP were observed at higher doses (50 and 100 µM) in hiPSC-vCMs which can be attributed to the heterogenous population, potential off-target effects at these high doses, as well as baseline expression of K_v_1.4 (I_to_).

Using nifedipine, we demonstrated the functional differences in Ca^2+^ handling dynamics between hiPSC-aCMs and -vCMs. Nifedipine elicited a dose-dependent response in hiPSC-vCMs demonstrating high sensitivity at 300 nM thereby confirming the functional presence of Ca_v_1.2. In contrast, hiPSC-aCMs were relatively insensitive to nifedipine showing no statistically significant differences in APD at all tested doses. This finding is further corroborated by the relatively decreased expression of *CACNA1C* (Ca_v_1.2) in the hiPSC-aCMs. This suggests that Ca^2+^ handling in hiPSC-aCMs may be reliant on other voltage-gated Ca^2+^ channels such as Ca_v_1.3, as this Ca^2+^ channel is blocked less potently by nifedipine^34^. Moreover, our qPCR assay confirmed that hiPSC-aCMs had higher expression of *CACNA1D* (Ca_v_1.3).

AVE0118 is an experimental K^+^ channel blocker (I_to_, I_Kur_, and I_Kr_) that was predicted to demonstrate targeted effects in hiPSC-aCMs. However, only a nuanced atrial specificity was observed in our assay. Although the effects were proportionally larger in hiPSC-aCMs, AVE0118 prolonged early repolarization of both hiPSC-aCMs and -vCMs in a similar fashion. The drug prolonged mid- and late-repolarization at a lower dose (1 µM) in hiPSC-aCMs showing minimal atrial specific effects. Interestingly, AVE0118 greatly affected Ca^2+^ handling in hiPSC-aCMs compared to hiPSC-vCMs with larger proportional prolongation of CaTD_50_ at all doses. These results were unexpected as AVE0118 is thought to be highly specific to hiPSC-aCMs due to its I_Kur_ blocking component. Perhaps the observed mixed-effects in both cell types is due to the drug binding to I_to_ (IC_50_: 3.4 µM) and I_Kr_ (IC_50_: 9.6 µM)^35^ which prolongs APD at the tested AVE0118 doses of 3 and 10 µM as genes encoding the channels producing the I_to_ (*KCNA4*) and I_Kr_ (*KCNH2*) were expressed in our hiPSC-vCMs. The drug has also been shown to be effective in terminating certain ventricular arrhythmias^36^ which was predicted based on our results of prolongation in the APD of hiPSC-vCMs.

Next, we used UCL1684, a highly specific SK channel pore blocker, to assess the presence of functional SK channels in hiPSC-aCMs. The SK channel has 3 paralogs but the SK3 channel variant (*KCNN3*) has been shown to be atrial-specific and has been implicated in AF pathogenesis in several GWAS studies^37,38^. In this study, UCL1684 displayed high specificity towards hiPSC-aCMs with a strong dose-dependent response. The drug confirmed the presence of functional SK channels in hiPSC-aCMs at 3 µM with a positive dose-dependent response while having no effect on hiPSC-vCMs at all tested doses (0.3, 1, 3, and 10 µM).

Vernakalant is touted as an atrial-selective compound clinically approved for intravenous cardioversion of AF^39^. Strikingly, out of all the tested drugs, vernakalant showed the most pronounced atrial-selective effects even though it is a blocker of multiple ion channels (I_Na_, I_Kur_, and I_K,Ach_). Vernakalant prolonged APD and CaTD of hiPSC-aCMs at three tested doses (3, 10, 30 µM). However, no statistically significant changes were observed in hiPSC-vCMs at early- and mid-repolarization while the slight prolongation at APD_80_ at the clinically relevant dose (30 µM) may be attributed to the I_Na_ blocking component of vernakalant. This result further demonstrates the sensitivity of the assay in establishing atrial-selective drug effects.

This study has several limitations. One limitation in our findings is that we cannot directly compare the results from qPCR and NanoString as both assays have fundamental differences in technical principles and statistical methodologies. Taken together, however, both assays show the global changes in cell type specific gene markers and further validate the role of retinoic acid in directing the cardiac differentiation process towards an atrial lineage. The main limitation in this field is the maturation state of the hiPSC-CMs as they have an overall immature phenotype with some crucial differences compared to adult cardiomyocytes^40^. Nonetheless, we were able to observe the stark differences in transcriptomic, protein, as well as functional signatures of AP and CaT in the two generated chamber-specific cell types. Additionally, maturation stage does not explain the differences in chamber-specific phenotype as parallel batch differentiation and time-in-culture were incorporated in our study design. Most importantly, we were able to capture effects of drugs that were expected to have atrial-specific properties in hiPSC-aCMs.

## Conclusion

The ability to differentiate hiPSC-aCMs provide a unique opportunity to study atrial physiology and its pharmacologic responses in a human-relevant *in vitro* model. We demonstrated an hiPSC-based *in vitro* model that recapitulates the molecular and functional characteristics of the phenotype of native atrial tissue. Our platform adds to the repertoire of cardiac drug screening and can be readily applied in future efforts of atrial-specific drug discovery.

## Acknowledgements

We would like to thank Ms. Salina Kung and Ms. Jennifer Yi for their help in designing the NanoString codeset. This work was financially supported by the Canadian Institutes of Health Research (GFT), the Canada Innovation Fund (GFT), the Stem Cell Network (GFT and ZL), and the Michael Smith Foundation (ZL).

## Data Availability Statement

Data can be made available upon reasonable request.

